# Cell to cell communication is an early event in glioblastoma

**DOI:** 10.1101/2020.01.24.917708

**Authors:** Marta Portela, Teresa Mitchell, Sergio Casas-Tintó

**Affiliations:** Instituto Cajal-CSIC. Av. del Doctor Arce, 37. 28002. Madrid. Spain; Department of Biochemistry and Genetics, La Trobe Institute for Molecular Sciences, La Trobe University, Melbourne, Australia

**Author notes:** Co-first authors (Marta Portela, Teresa Mitchell).

**Keywords:** Glia, Cancer, Glioblastoma, Tumour Microtubes, JNK, Neurodegeneration

## Abstract

Glioblastoma (GB) is the most aggressive and lethal tumour of the central nervous system (CNS). GB cells proliferate rapidly and display a network of ultra-long tumour microtubes (TMs) that mediate cell to cell communication. GB TMs infiltrate into the brain, enwrap neurons and facilitate the depletion of Wingless (Wg)/WNT from the neighbouring neurons. GB cells establish a positive feedback loop including Wg signalling upregulation that activates the JNK pathway and matrix metalloproteases (MMPs), in turn, these signals promote TMs infiltration, GB progression and neuronal synapse loss and degeneration. Thus, cellular and molecular signals other than primary mutations emerge as central players of GB. Here we describe the temporal organization of the events that occur in GB. We define the progressive activation of JNK pathway signalling mediated by Grindelwald (Grnd) receptor, is caused by the ligand Eiger (Egr)/TNFα produced by the healthy tissue. We propose that cellular interactions of GB with the rest of the brain is an early event that precedes GB proliferation and expansion. We conclude that non-autonomous signals facilitate GB progression and contribute to the complexity and versatility of these incurable tumours.

## Introduction

Glioblastoma multiforme (GB) is the most frequent and aggressive primary malignant brain tumour with a 3 per 100.000 incidence per year (Gallego, 2015). GB patients’ median survival is 12-15 months, with less than 5% of survival after 5 years (Gallego, 2015; Louis et al., 2016; McGuire, 2016; Rogers et al., 2018). The causes of GB are under debate (McGuire, 2016), 5% of the patients develop GB after a low grade astrocytoma (Alifieris and Trafalis, 2015) and the most frequent mutations include gain of function of the epidermal growth factor receptor (EGFR) (97%) and the phosphatidylinositol-3 kinase (PI3K)/phosphatase and tensin homologue (PTEN) pathways (88%) (Hayden, 2010). The diagnosis, and therefore the treatment of GB, requires a mutation analysis considering the high frequency of clones within the same primary GB (Wick et al., 2018). Temozolomide (TMZ) is the only treatment for GB, however, recent discoveries restrict the use of TMZ in GB patients depending on the methylation status of methylguanine DNA methyltransferase (MGMT) *(Wick et al., 2018)*. Moreover, among other mutations, *Isocitrate dehydrogenase (IDH)* define the nature and features of GB (Waitkus et al., 2018) together with molecular alterations including 1p/10q deletions and *tumour suppressor protein 53 (TP53)* and *alpha thalassemia/mental retardation (ATRX)* mutations (Miller et al., 2017; Waitkus et al., 2018). The genetic and molecular heterogeneity complicate the diagnosis and treatment of these fatal brain tumours.

EGFR mutant forms show constitutive kinase activity that chronically stimulates Ras signalling to drive cell proliferation and migration (Furnari et al., 2007; Maher et al., 2001). Other common genetic lesions include loss of the lipid phosphatase PTEN, which antagonizes the phosphatidylinositol-3 kinase (PI3K) signalling pathway, and mutations that activate PI3KCA, which encodes the p110a catalytic subunit of PI3K (Furnari et al., 2007; Maher et al., 2001). GBs often show constitutively active Akt, a major PI3K effector. Multiple mutations that coactivate EGFR-Ras and PI3K/Akt pathways are required to induce a glioma (Holland, 2000). In *Drosophila*, a combination of EGFR and PI3K mutations effectively causes a glioma-like condition that shows features of human gliomas including glia expansion, brain invasion, neuron dysfunction, synapse loss and neurodegeneration This model involves the co-overexpression of constitutively active forms of Epidermal Growth Factor Receptor (dEGFR^λ^) and an activated membrane-localized version of the PI3K catalytic subunit p110α/PI3K92E (dPI3K92E^CAAX^) under Gal4 UAS control, specifically driven in the glial cells by means of *repo-Gal4* (Brand and Perrimon, 1993; Casas-Tinto et al., 2017).

The recent discovery of a network of ultra-long tumour microtubes (TMs) in GB (Osswald et al., 2015), also known as cytonemes in *Drosophila* (Casas-Tinto and Portela, 2019), improves our understanding of GB progression and therapy resistance (Osswald et al., 2016). In GB, glial cells display a network of membrane projections which mediate cell to cell communication. TMs are actin-based filopodia that infiltrate into the brain and reach long distances within the brain (Osswald et al., 2015). TMs are required in GB cells to mediate Wingless (Wg)/WNT signalling imbalance among neurons and GB cells. Wg/WNT signalling is increased in GB cells to promote proliferation, at the expense of neuronal Wg signalling that results in neurodegeneration and lethality (Arnes and Casas Tinto, 2017; Casas-Tinto and Portela, 2019; Portela et al., 2019). The central role of TMs in GB biology has emerged as a fundamental mechanism for GB rapid and lethal progression; thus, it is an attractive field of study towards potential GB treatments. However, the molecular mechanisms underlying the expansion of TMs and the signalling pathways mediating TM infiltration are still poorly understood.

The Jun-N-terminal Kinase (JNK) pathway is a hallmark of GB cells that is associated to glial proliferation and stem-like status, and currently it is a pharmacological target for GB (Matsuda et al., 2012). Moreover, the JNK pathway is the main regulator of Matrix metalloproteases (MMPs) expression and cell motility in many organisms and tissues including tumors like GB (Cheng et al., 2012; Ispanovic and Haas, 2006; Lee et al., 2009; Portela et al., 2019; Uhlirova and Bohmann, 2006; Zeigler et al., 1999).

MMPs are a family of endopeptidases capable of degrading the extracellular matrix (ECM). Members of the MMP family include the “classical” MMPs, the membrane-bound MMPs (MT-MMPs), the ADAMs (a disintegrin and metalloproteinase; adamlysins) and the ADAMTS (a disintegrin and metalloproteinase with thrombospondin motif). There are more than 20 members in the MMP and ADAMTS family including the collagenases, gelatinases, stromelysins, some elastases and aggrecanases (Malemud, 2006). The vertebrate MMPs have genetic redundancy and compensation, they have overlapping substrates, and pharmacological inhibitors are non-specific. There are two orthologues to human MMPs in *Drosophila, MMP1* and *MMP2*. MMP1 is secreted and MMP2 is membrane-anchored (Page-McCaw et al., 2003). However, recent reports propose that products of both genes are found at the cell surface and released into media, and that GPI-anchored MMPs promote cell adhesion when they become inactive. Moreover, the two MMPs cleave different substrates, suggesting that this is the important distinction within this small MMP family (LaFever et al., 2017). MMPs are upregulated in several tumours, including GBs. Cancer cells produce MMPs to facilitate tumour progression and invasiveness and MMPs upregulation in GB is associated with the diffuse infiltrative growth and have been proposed to play a role in GB cell migration and infiltration (de Lucas et al., 2016; Veeravalli and Rao, 2012) reviewed in (Nakada et al., 2003). MMPs are upregulated in human GB cell-lines and biopsies as compared with low-grade astrocytoma (LGA) and normal brain samples (Hagemann et al., 2012; Hagemann et al., 2010). In particular, among the 23 MMPs present in humans, MMP9, MMP2 and MMP14 are directly implicated in growth and invasion of GB cells (Munaut et al., 2003).

WNT induces MMPs expression during development and cancer (Lowy et al., 2006; Lyu and Joo, 2005; Page-McCaw et al., 2003; Roomi et al., 2017; Uraguchi et al., 2004) associated to cell migration and metastasis. Specifically, in human GB, MMP2 expression and their infiltrative properties correlate with Wnt5 (Kamino et al., 2011; Roth et al., 2000) and MMP9 is upregulated upon EGFR activity (Chen et al., 2016).

In consequence, MMPs upregulation in GB is an indicator of poor prognosis (Yamamoto et al., 2002) and the study of the mechanisms mediated by MMPs is relevant for the biology of GB, and cancer in general. GB cells project TMs which cross the extracellular matrix (ECM) and infiltrate in the brain to reach territories distant from the primary GB site (Osswald et al., 2015; Osswald et al., 2016). We have previously demonstrated that GB cells activate the JNK pathway and accumulate MMPs (MMP1 and 2). MMPs contribute to TMs expansion in the brain and facilitate Frizzled1-mediated Wg/WNT signalling in GB cells. Moreover, Wg/WNT signalling mediates JNK activation in GB cells to continue with the TMs infiltration process. We hypothesize that the founder mutations in GB (PI3K and EGFR) initiate the process with the expansion of the TMs; afterwards, the system self-perpetuates (TMs-Fz1/Wg-JNK-MMPs-TMs) to facilitate GB progression and infiltration in the brain (Portela et al., 2019).

We have recently showed that JNK signalling pathway is required for GB progression and proliferation. JNK pathway activation mediated by the receptor Grnd is a requirement for both TMs expansion, and Fz1-mediated Wg depletion from neurons. In turn, Wg pathway upregulation in GB induces JNK activity in GB cells that mediate the production of matrix metalloproteinases (MMPs), moreover *MMPs* silencing in GB cells is sufficient to rescue neurodegeneration and premature death caused by GB (Portela et al., 2019). However, the molecular mechanisms that determine JNK pathway activation in GB cells remain unknown.

Here, we describe the role of the ligand Egr in the activation of JNK pathway in GB through the specific receptor Grnd, highlighting again the contribution of communication signals between healthy tissue (neurons) and GB cells to the progression of the disease (Jarabo et al., 2020; Portela et al., 2019). Egr is expressed and secreted by non-tumoral tissue, but it is accumulated in tumoral cells and activates the JNK pathway. In consequence, GB cells produce MMPs that facilitate TM progression and GB dissemination. At the cellular level, we analyse 3 aspects required for GB progression: TMs expansion, GB proliferation and synapse loss in the surrounding neurons, and propose a timeline for these events in GB progression.

## RESULTS

### Combined activation of EGFR and PI3K pathways is required for GB progression

The *Drosophila* GB model reproduces the main features of the disease including the expansion of TMs. Cytonemes and TMs share multiple features (Casas-Tinto and Portela, 2019; Portela et al., 2019) and can be visualized with membrane tags (e.g., CD8:GFP or myr-RFP), or cytoneme components including signalling proteins such as Ihog (Ihog-RFP). Membrane markers are expressed under the control of the glial specific enhancer *repo-Gal4* (Casas-Tinto et al., 2017). Therefore, only glial cells are marked with RFP, and in the *Drosophila* brain the unlabeled surrounding cells are neuroblasts and neurons (Figure S1A) (Portela et al., 2019).

To determine the separate contribution of PI3K and EGFR signaling pathways to GB progression, we first analyzed EGFR and PI3K/PTEN pathways independently. Fz1 localization in the TMs is key to trigger Wg signaling and GB proliferation. We assessed the presence of Fz1 receptor in glial membranes with a specific monoclonal antibody previously validated (Portela et al., 2019). Fz1 receptors signal is localized homogeneously across the brain in control samples (Figure S1B). However, in the GB model (PI3K+EGFR) Fz1 accumulate in TMs (Figure S1C (Portela et al., 2019)). To determine if expression of constitutively active forms of PI3K or EGFR independently is sufficient to trigger TMs network expansion and Fz1 localization in the TMs, we expressed *Drosophila PI3K* or *EGFR* (*UAS-dp110*^*CAAX*^ *or UAS-TOR-DER*^*CA*^) in glial cells driven by *repo-Gal4* and stained with Fz1 antibody. The results show that Fz1 receptors localized homogeneously across the brain (Figure S1D-E) similar to control samples, and the glial TM network does not expand (compare Figure S1D-E to Figure S1B-C). These data suggest that the activation of both pathways together is necessary for the expansion of the TMs network and Fz1 localization in the TMs.

PI3K and EGFR pathways converge in *dMyc* expression and *dMyc* expression is required for GB development (Annibali et al., 2014; Read et al., 2009; Tateishi et al., 2016; Wang et al., 2017). Thus, to determine if d*Myc* is sufficient to trigger TMs network and Fz1 accumulation, we ectopically overexpressed d*Myc* (*UAS-dMyc*) in glial cells under *repo-Gal4* regulation and stained the brains with Fz1 (Figure S1F), Wg (Figure S1G-I) and Cyt-Arm antibodies (Figure S1J-L). The confocal images show no morphological evidence of TMs expansion or GB formation (Figure S1F compare to glioma in S1C). Moreover, Fz1, Wg or Cyt-Arm protein distribution resemble control brains (Figure S1B, F, G, I, J, L). Taking these results together, *dMyc* overexpression is not sufficient to reproduce the features of the GB even though it is a convergent point of EGFR and PI3K pathways. These results suggest that both PI3K and EGFR constitutive upregulation in glial cells is necessary for the expansion of glial TMs, Fz1 localization in TMs and activation of the Wg pathway in glial transformed cells.

Next, to determine the epistatic relations behind *MMPs* upregulation, we used a specific monoclonal antibody to visualize MMP1. The confocal images show that MMP1 is homogeneously distributed across the brain in control samples (Figure 1A) and accumulates in TMs in GB samples (Figure 1B). Besides, MMP1 shows a homogeneous distribution through the brain upon constitutive activation of PI3K (*dp110*^*CAAX*^) and EGFR (*TOR-DER*^*CA*^) in glial cells (Figure 1C-D, quantified in F). Besides, *dMyc* overexpression in glial cells did not cause significant changes in MMPs localization (Figure 1E-F). Taking these results together, independent overexpression of EGFR, PI3K or dMyc is not sufficient to reproduce the features of GB. These results suggest that the combined activity of PI3K and EGFR pathways are necessary to activate a downstream pathway responsible for the expansion of TM glial projections and MMP1 accumulation in GB cells, however *dMyc* overexpression is not sufficient to cause these phenotypes.

**Figure 1:**
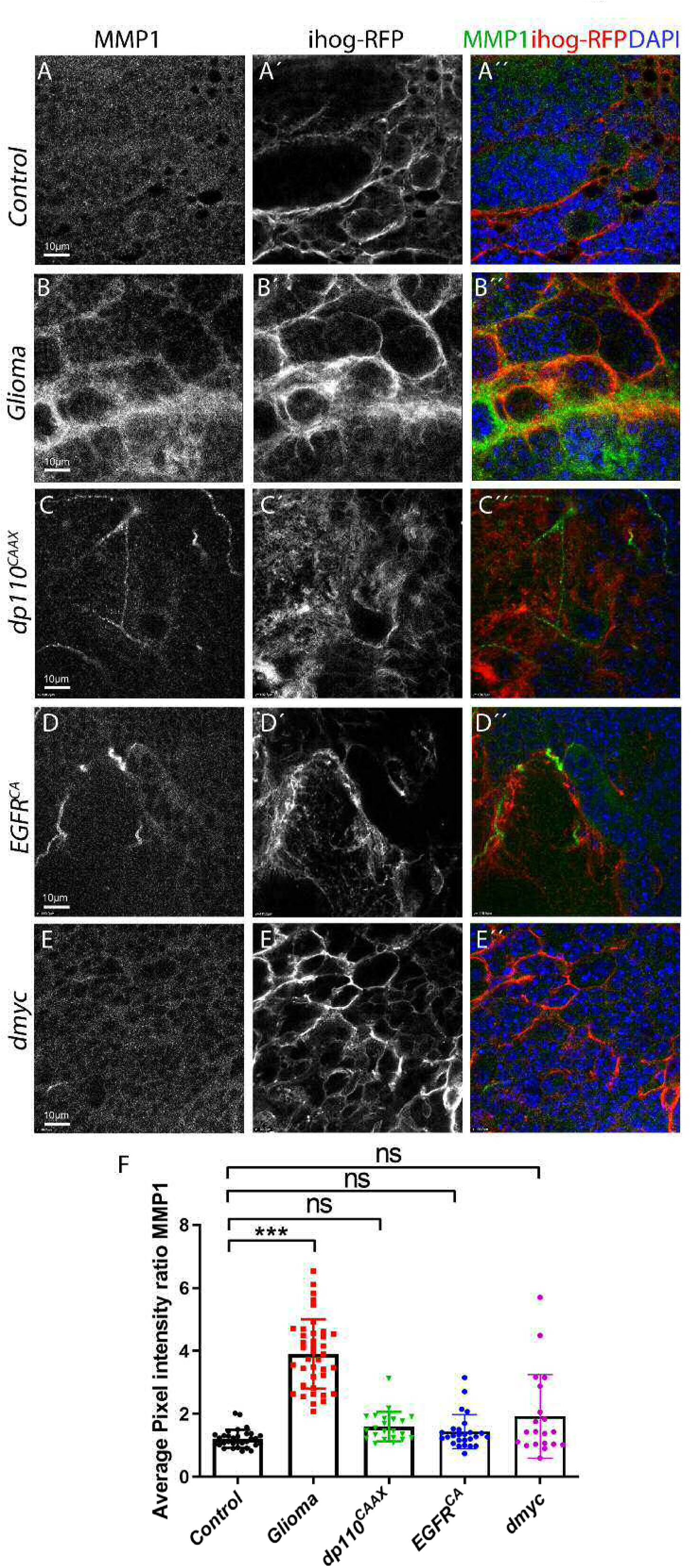
Independent constitutive activation of PI3K or EGFR or ectopic *dmyc* are not responsible for MMP1 accumulation in GB. Brains from 3rd instar larvae. Glia are labeled with *UAS-Ihog-RFP* driven by *repo-Gal4* to visualize active cytonemes/ TM structures in glial cells and stained with MMP1 (green). (A) MMP1 (green) is homogeneously distributed in control. (B) In Glioma brains, MMP1 accumulates in the TMs and specifically in the TM projections that are in contact with the neuronal clusters. (C-E) MMP1 (green) is homogeneously distributed in (B) *dp110*^*CAAX*^, (C)*TOR-DER*^*CA*^ and (D) *dmyc* sections. The glial cytonemes (red) revealed by Ihog-RFP do not overgrow or encapsulate neuronal clusters. Nuclei are marked with DAPI (blue). (F) Quantification of MMP1 staining ratio between iHog^+^ and iHog^−^ domains. Kruskal– Wallis test with Dunns post-test. Error bars show S.D. *** P<0.001 or ns for non-significant. Scale bar size is indicated in this and all figures. Genotypes: (A) *UAS-lacZ/repo-Gal4, UAS-ihog-RFP* (B) *UAS-dEGFR*^*λ*^, *UAS-dp110*^*CAAX*^;; *repo-Gal4, UAS-ihog-RFP* (C) *UAS-dp110*^*CAAX*^;; *repo-Gal4, UAS-ihog-RFP* (D) *UAS-TOR-DER*^*CA*^;; *repo-Gal4, UAS-ihog-RFP* (E) *repo-Gal4, UAS-ihog-RFP/UAS-dmyc*

### Non-autonomous activation of JNK pathway in GB

In addition to Wg pathway, GB cells activate JNK pathway to maintain the stem-like GB cells, which has become a pharmacological target for the treatment of GB (Matsuda et al., 2012). A recent study using gene expression profiling identified genes that were significantly correlated with the overall survival in patients with GB (Hsu et al., 2019). 104 genes were identified, which are common between patients with GB and those with low grade gliomas and can be used as core genes related to patient survival. Of these, 10 genes (CTSZ, EFEMP2, ITGA5, KDELR2, MDK, MICALL2, MAP 2 K3, PLAUR, SERPINE1, and SOCS3) can potentially classify patients with gliomas into different risk groups. Among these pathways, the TNF-alpha signalling pathway stands out. Four genes from this 10-gene group (MAP 2 K3, PLAUR, SERPINE1, and SOCS3) are involved in TNFα signalling, and they might have potential prognostic value for patients with GB (Hsu et al., 2019).

TNF-like weak inducer of apoptosis (TWEAK, also known as TNFSF12) is one of the ligands of the tumor necrosis factor (TNF) family (Chicheportiche et al., 1997). TNFSF12 preferentially activates non-canonical NF-κB and promotes the invasive properties of glioma cells (Cherry et al., 2015). To determine the contribution of TNFSF12 to GB, we analysed the data from the CGGA dataset from GlioVis (http://gliovis.bioinfo.cnio.es/). In silico analysis showed that TNFSF12 gene is highly upregulated in several samples of CNS tumors, including GB (Figure 2A). Accordingly, *TNFSF12* expression has a negative implication in the overall survival of patients with gliomas (Figure 2B).

**Figure 2:**
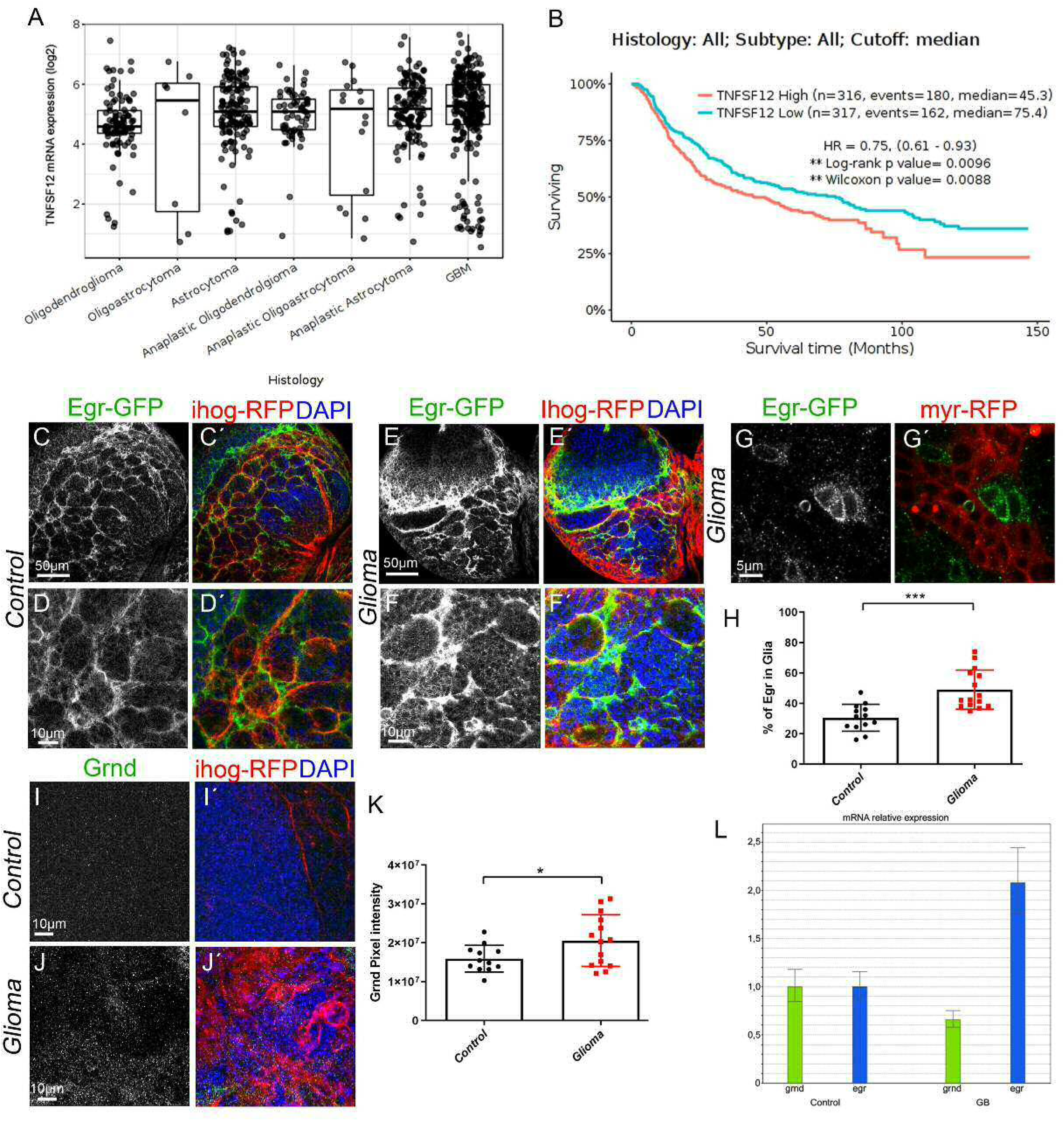
Egr re-localizes from neuron to glia and its receptor Grindelwald is accumulated in GB. (A-B) In silico analysis of the data from the CGGA dataset from GlioVis (http://gliovis.bioinfo.cnio.es/) showing that TNFSF12 is highly upregulated in several tumors of the CNS of glial origin, including GB (A). In silico analysis of CGGA data indicating the survival of patients with high or low TNFSF12 expression (B). Brains from 3rd instar larvae. Glia are labelled with *UAS-Ihog-RFP* driven by *repo-Gal4* to visualize active cytonemes/ TM structures in glial cells, carry a Egr-GFP reporter (green). (C-D) In control brain sections Egr-GFP (green) signal is localized mostly (70%) in neurons in close contact with glial cells. (E-G) In glioma brain sections Egr-GFP (green) signal shifts and it is 50% localized in glioma cells. Higher magnification shown in (G) (H) Quantification of the percentage of Egr in glia in control and glioma samples. (I-J) Grnd staining (green) in control and glioma samples. (K) Quantification of the Grnd pixel intensity. Nuclei are marked with DAPI (blue). (L) qPCRs with RNA extracted from control and glioma larvae showing upregulation of the transcription (mRNA levels) of *egr*. Two-tailed t test with Welch-correction. Error bars show S.D. * P<0.05, ***p≤0.001. Scale bar size are indicated in this and all figures. Genotypes: (C-D) Egr-GFP; *repo-Gal4, ihog-RFP/UAS-lacZ* (E-F) *UAS-dEGFR*^*λ*^, *UAS-dp110*^*CAAX*^;*Egr-GFP; repo-Gal4, UAS-ihog-RFP* (G) *Gal80*^*ts*^*/UAS-dEGFR*^*λ*^, *UAS-dp110*^*CAAX*^; *Egr-GFP; repo-Gal4, myr-RFP* (I) *repo-Gal4, ihog-RFP/UAS-lacZ* (J) *UAS-dEGFR*^*λ*^, *UAS-dp110*^*CAAX*^;; *repo-Gal4, UAS-ihog-RFP*

Due to the relevance of JNK pathway and TNFα ligand in GB prognosis we decided to study Egr, that is the solely *Drosophila* orthologue of the ligand for the mammalian Tumour Necrosis Factor (TNF)-TNF Receptor signalling pathway (Igaki et al., 2002; Moreno et al., 2002) and its receptor Grnd (Andersen et al., 2015), in the *Drosophila* GB model.

### Egr re-localizes from Neuron to Glia in GB

To monitor Egr protein distribution, we used a transgenic *Drosophila* line in which the endogenous *egr* gene is GFP tagged (Egr-GFP protein fusion) and monitored GFP signal. The results from confocal microscopy show that 70% of Egr-GFP signal is localized in the neurons that are in the proximity of glial cells and 30% of Egr-GFP signal is localized in the glia (Figure 2C-D, H). However, in GB samples there is a shift of Egr-GFP signal from neurons to glia (∼50%) (Figure 2E-H). Next, we monitored the expression pattern of the JNK receptor Grnd in GB and control samples, with a specific antibody previously validated (Andersen et al., 2015). The quantification of Grnd signal in glial membranes show an increase of Grnd protein in GB brains (Figure 2I-K). The abnormal distribution of Egr and Grnd in glioma brains could be due to either an increase in gene and/or protein expression or to redistribution of the proteins. We undertook quantitative Polimerase Chain Reaction (qPCRs) experiments with RNA extracted from control and GB larvae brains, which revealed an that there is a 2-fold increase of *egr* transcription in GB brains (Figure 2L) consistent with the increased Egr staining in glioma brains.

### Egr expressed in healthy tissue activates JNK in GB

Next we studied which cell type upregulates *egr* expression in GB brains. To determine whether the source of Egr are the glial cells, we silenced *egr* expression specifically in glioma cells. We used two different RNAi lines to knockdown *egr* (*egr-RNAi*) in GB cells (*repo>dp110*^*CAAX*^; *EGFR*^*λ*^; *egr-RNAi*). The quantifications show that *egr* knockdown does not prevent GB cell proliferation (Fig 3A–3D and 3E) nor GB membrane expansion (Fig 3A-3D and 3F). These results suggest that *egr* expression in GB cells is not relevant for GB progression in the *Drosophila* model. Therefore, taking into consideration that there is an increase of Egr protein in glial membranes in GB compared to control brains (Figure 2), and it is not due to a transcriptional upregulation of *egr* specifically in the glial cells, we propose that Egr is transcriptionally upregulated specifically in neurons and re- localised from the surrounding neuronal tissue to GB cells in order to activate JNK pathway through Grnd receptor in GB cells.

**Figure 3:**
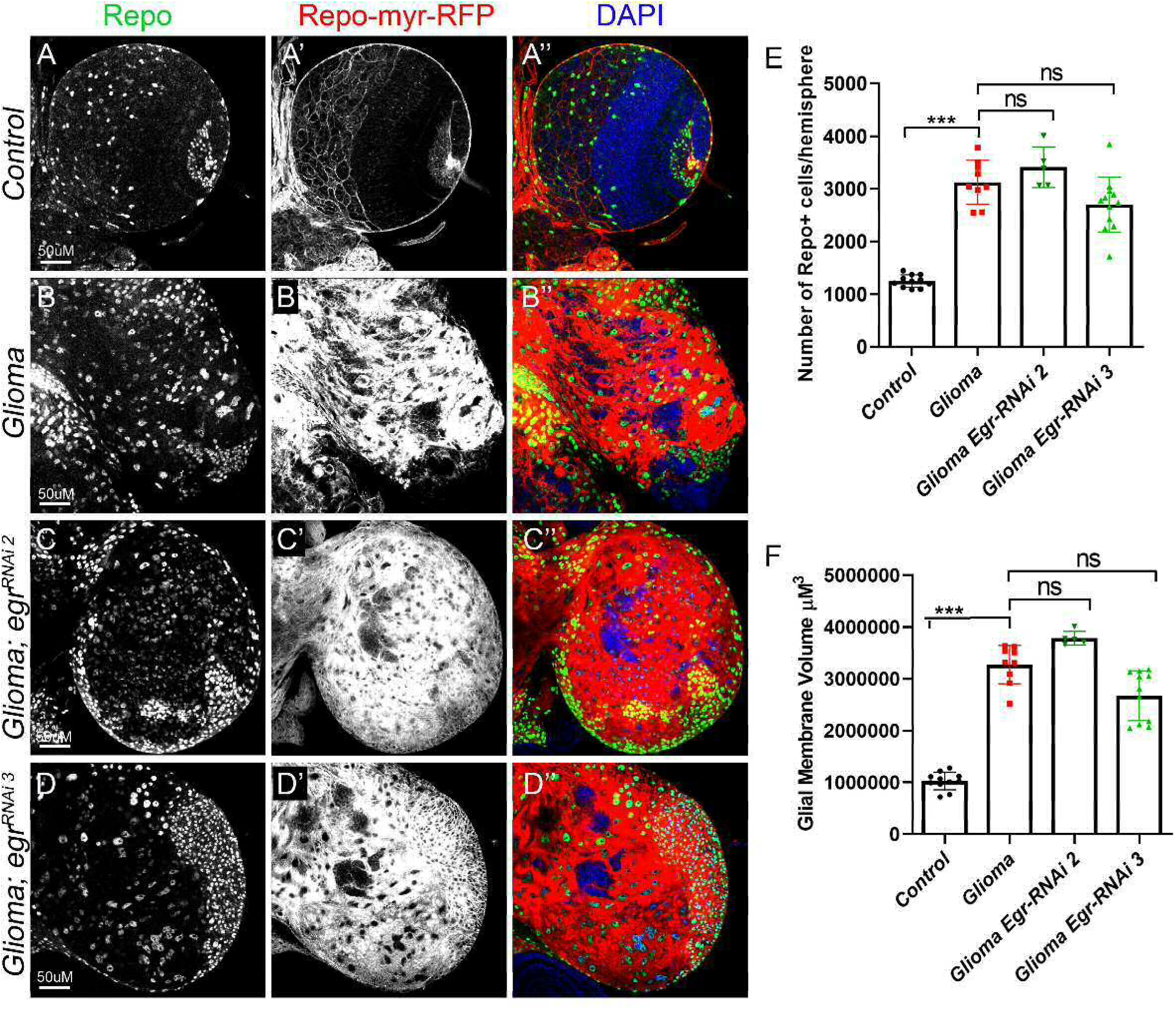
Egr expressed in healthy tissue activates JNK in GB. Egr expression in glioma cells is dispensable for GB progression. (A–D) Larval brain sections with glial membrane projections labelled in grey (red in the merge) and glial cell nuclei stained with Repo (grey, green in the merge). (C-D) *egr* knockdown in GB cells (*egr-RNAi*) does not prevent GB cell number increase nor GB TM volume expansion, quantified in (E–F). One-way ANOVA with Bonferroni post-test (E) and Kruskal–Wallis test with Dunns post-test (F). Error bars show S.D. *** P<0.001 or ns for non-significant. Genotypes: (A) *Gal80*^*ts*^*/repo-Gal4, myr-RFP/UAS-lacZ* (B) *Gal80*^*ts*^*/UAS-dEGFR*^*λ*^, *UAS-dp110*^*CAAX*^;; *repo-Gal4, myr-RFP* (C) *Gal80*^*ts*^*/UAS-dEGFR*^*λ*^, *UAS-dp110*^*CAAX*^; *UAS-egr-RNAi; repo-Gal4, myr-RFP* (D) *Gal80*^*ts*^*/UAS-dEGFR*^*λ*^, *UAS-dp110*^*CAAX*^;; *repo-Gal4, myr-RFP/ UAS-egr-RNAi*

### Progressive JNK activation in glioma

JNK is upregulated in several tumours including GB and it is related to glioma malignancy (Hagemann et al., 2005; Huang et al., 2003; Mu et al., 2018; Zeng et al., 2018). Moreover, JNK is a target for specific drugs in combination with temozolomide treatments as it was proven to play a central role in GB progression (Feng et al., 2016; Kitanaka et al., 2013; Matsuda et al., 2012; Okada et al., 2014). However, little is known about the molecular mechanisms underlying JNK activation in GB cells and the functional consequences for GB progression.

We had previously confirmed JNK pathway activation in *Drosophila* GB cells (Portela et al., 2019), by using the *TRE-RFP* reporter that confer transcriptional activation in response to JNK signalling (Chatterjee and Bohmann, 2012; Jemc et al., 2012; Ruan et al., 2016). We decided to study the temporal activation of the JNK pathway to uncover the order the signalling events that occur in GB cells and healthy surrounding tissue. We took advantage of the JNK reporter (*puc-LacZ*) that monitors the transcriptional activation of the downstream JNK target *puckered* (Langen et al., 2013; Martin-Blanco et al., 1998) to analyse JNK activity in GB at 48 and 96h after GB induction. To control the temporal induction of the tumour, we used the thermo sensitive repression system Gal80^TS^. Individuals maintained at 17°C did not activate the expression of the UAS constructs, after larvae were switched to 29°C, the protein Gal80^TS^ was not longer able to repress Gal4 system and the genetic induction of GB was activated. In addition, GB cells were marked with RFP (*UAS-myr-RFP*) to distinguish GB cells and healthy tissue.

We quantified the bGalactosidase (LacZ) signal that overlaps with RFP. The analysis of confocal images showed that puc*-LacZ* is mostly activated in neurons (∼78%) in control brains. 48h after the induction of the GB, *puc-LacZ* activation in neurons is reduced (∼37%) and GB cells show a progressive activation of JNK in GB cells: from 63% of *puc-LacZ* signal in glia 48h after the tumour induction, and ∼80% 96h after tumour induction (Figure 4A-D), indicating that JNK pathway is progressively activated in GB cells. These results indicate that JNK pathway activation increases in GB cell as the tumour progresses.

**Figure 4:**
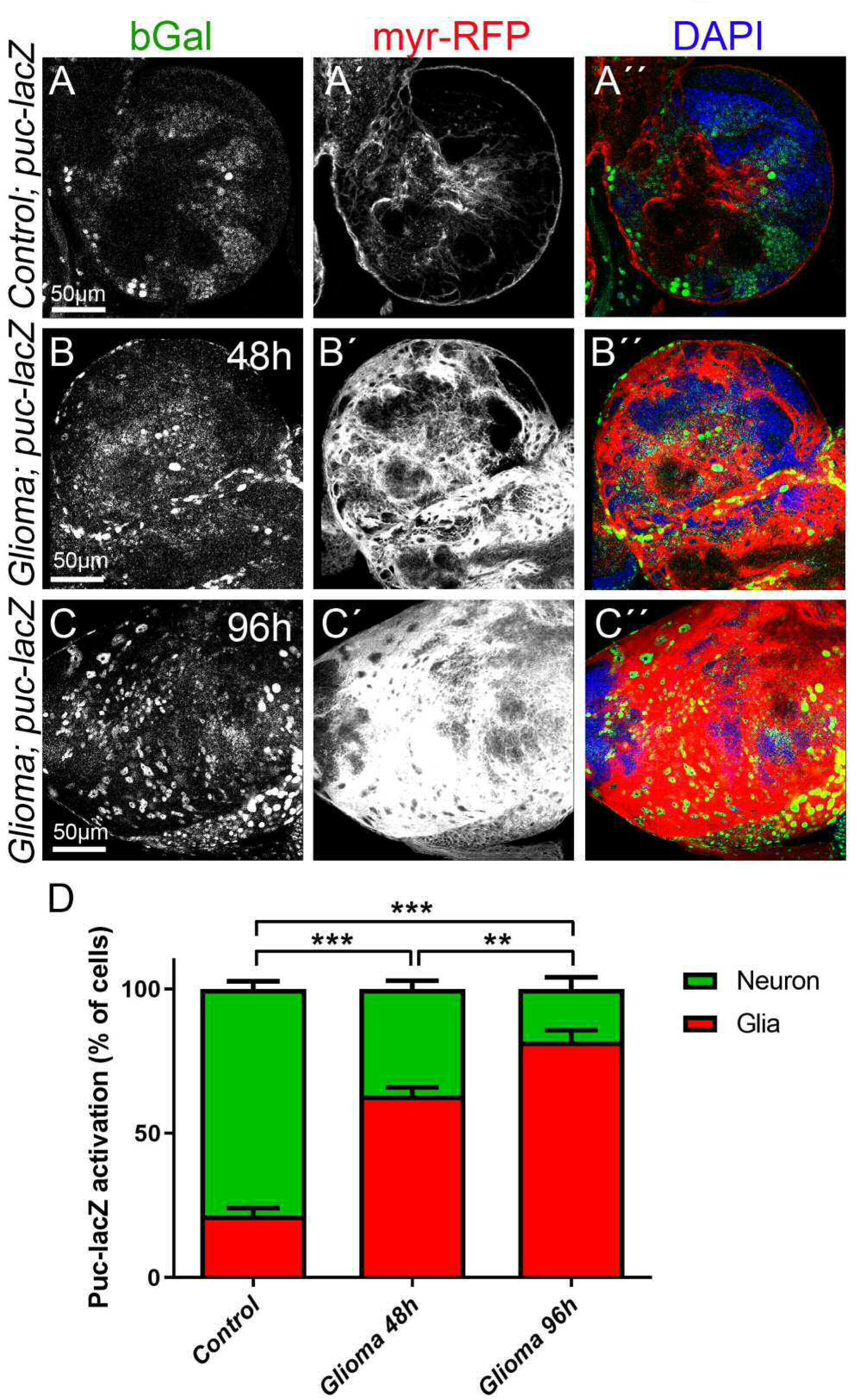
JNK signalling pathway activation in GB. Larval brain sections from 3rd instar larvae displayed at the same scale. Glial cell bodies and membranes are labelled with *UAS-myr-RFP* (red) driven by *repo-Gal4* (A-C) JNK signalling pathway reporter *puc-lacZ* in control, glioma 48h and glioma 96h after tumour induction. *puc-lacZ* shows activation of the pathway mostly in neurons in control samples (A) and then shows a progressive activation in GB cells (B-C). (D) Quantification of the % of cells with *puc-lacZ* activation in glial cells and neurons. Nuclei are marked with DAPI (blue). One-way ANOVA with Bonferroni post-test. Error bars show S.D. ** P<0.01, *** P<0.001. Scale bar size is indicated in this and all figures. Genotypes: (A) *Gal80*^*ts*^*/repo-Gal4, myr-RFP/puc-lacZ* (B-C) *Gal80*^*ts*^*/UAS-dEGFR*^*λ*^, *UAS-dp110*^*CAAX*^;; *repo-Gal4, myr-RFP/puc-lacZ*

### Timeline: GB first causes neurodegeneration, then TMs infiltration and cell number increase

The previous results indicate that JNK signalling pathway activation is progressive, and therefore GB progression follows a chronogram. To evaluate the cellular events associated to GB (progressive growth of the glial membrane projections and the number of glial cells), we dissected larval brains at 24, 48 and 72h after tumour induction (referred to as day 1D, 2D or 3 days in Figure 5A-C). GB cells membrane was visualized with *UAS-myr-RFP* and we stained the glia nuclei with a specific anti-Repo antibody to quantify the glial cell number by analysing confocal microscopy images from entire brains (Figure 5A-C).

**Figure 5.**
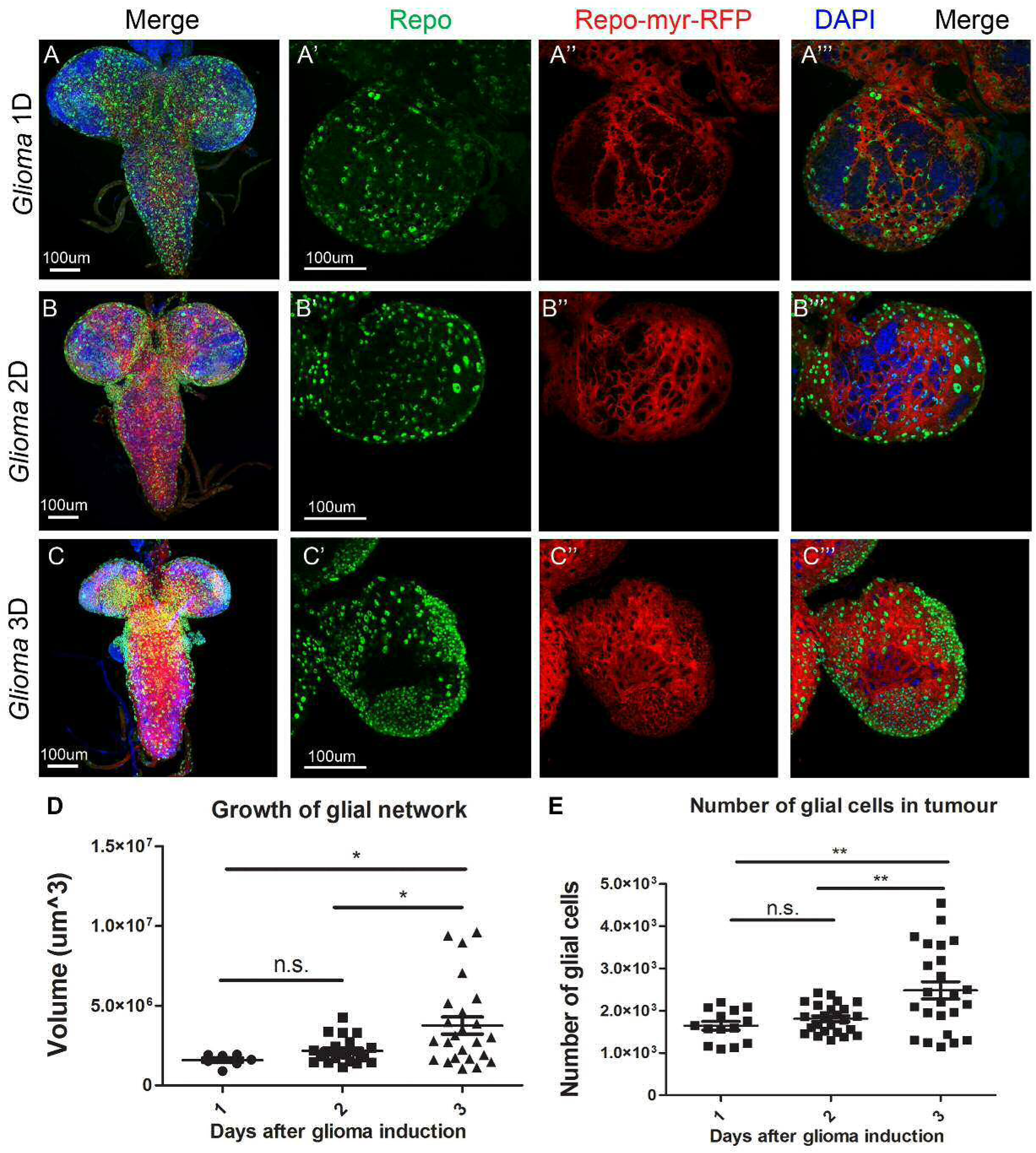
GB TM network volume and number of glial cells 1, 2 and 3 days after tumor induction. Third-instar larval brains or optical lobes Repo+ cells (glial cells) are marked with anti- Repo and visualized as green spots. TMs are marked in red via *UAS-myrRFP* which labels the plasma membrane of glial cells with RFP. (A) Third-instar larval brain 1 day after tumor induction. (B) Third-instar larval brain 2 days after tumor induction. (C) Third- instar larval brain 3 days after tumor induction. (D) Graphical representation of the TM volume at the different timepoints after tumor induction. (E) Graphical representation of the number of glial cells in brains at the different timepoints after tumor induction. Nuclei are marked with DAPI (blue). One-way ANOVA with Bonferroni post-test. *p≤0.05 **p≤0.01, n.s.=not significant. Genotypes: (A-C) *UAS-dEGFR*^*λ*^, *UAS-dp110*^*CAAX*^; *Gal80*^*ts*^; *repo-Gal4, myr-RFP*

The statistical analysis of glial membrane volume quantifications (Figure 5D) shows no significant increase in the volume of the network between day 1 and day 2 after tumor induction. Nevertheless, there is a significant increase in the volume of the network between day 2 and day 3 after tumor induction. Similarly, the statistical analysis of the number of glial cells (Figure 5E) shows no significant increase in the number of glial cells between day 1 and day 2, but there is a significant increase between day 2 and day 3, and between day 1 and day 3 after tumor induction. This suggests a progressive growth and expansion of the glial membranes and a progressive increase in the number of GB cells.

To evaluate the impact of the progressive GB proliferation, growth and membrane expansion on the surrounding neurons, we studied the neurons. Motor neurons are localized in the CNS and project their axons towards the neuromuscular junction (NMJ), a well-established system to evaluate neurodegeneration (Keshishian et al., 1996; Portela et al., 2019).

We quantified the number of synapses in the NMJ of third-instar larvae 1, 2 and 3 days after GB induction. NMJs were stained with anti-bruchpilot to visualize active zones (synapses) by using confocal microscopy (Figure 6A-C).

**Figure 6.**
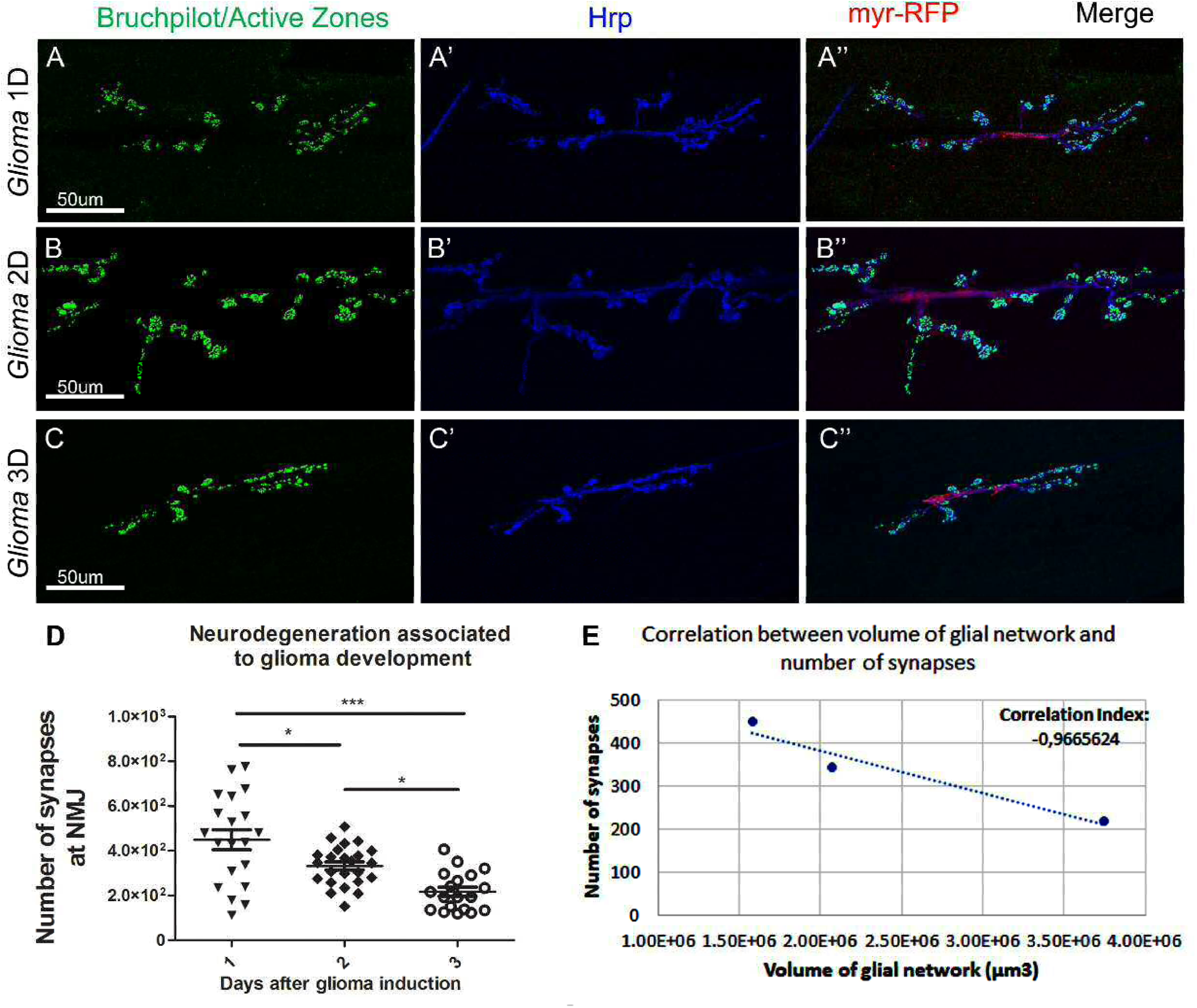
Number of synapses at the NMJ of third-instar larvae 1, 2 and 3 days after tumor induction. The synaptic connexions at the NMJ of *Drosophila* larvae are between muscles 6 and 7 in the third and fourth abdominal segment. Anti-Bruchpilot (α-NC82) binds to Bruchpilot marking presynaptic zones in green. Anti-Hrp marks the neuron membrane in blue. Glial cell membranes are visualized in red via *UAS-myrRFP*. (A) Third-instar larval NMJ 1 day after tumor induction. (B) Third-instar larval NMJ 2 days after tumor induction. (C) Third-instar larval NMJ 3 days after tumor induction. (D) Graphical representation of the number of synapses at the NMJ 1, 2 and 3 days after tumor induction. One-way ANOVA with Bonferroni post-test. ***p≤0.001, * p≤0.05. (E) Scatter plot showing the correlation between volume of the TMs and number of synapses at the NMJ. There is a negative correlation between the volume of the network and the number of synapses, therefore, at smaller volumes of glial network there are more synaptic connexions at the NMJ and as the volume increases, more synapses are lost. Genotypes: (A-C) *UAS-dEGFR*^*λ*^, *UAS-dp110*^*CAAX*^; *Gal80*^*ts*^; *repo-Gal4, myr-RFP*

The statistical analysis of synapse number (Figure 6D) show progressive synapse loss between day 1 and day 2, and between day 2 and day 3 after tumor induction. The overall loss of synapses at the NMJ is highly significant (Figure 6D).

To determine whether there is an association between the volume of the glial network and the number of synapses at the NMJ, we plotted the number of synapses against the volume of the glial network in a correlation graph (Figure 6E). The correlation index is -0.966 (3s.f.) indicating there is a negative association between the volume of the TMs and the number of synapses at the NMJ. Therefore, larger glial membrane network leads to a greater synapse loss. Taking all these results together, the first cellular event happening after the tumor induction is the neurodegeneration caused by GB, suggesting that the interaction between GB cells and neurons is an early event. Then we observed the expansion of the TMs network volume and finally, the increase in the number of glial cells. These results suggest that the interaction and activation of signaling pathways between GB cells and neurons occurs very early in the process of GB progression, bringing the study of cell to cell communication as an intriguing feature of GB.

## Discussion

Activating mutations for EGFR and PI3K pathways are the most frequent initial signals in GB. However, the attempts to treat GB reducing the activation of these pathways have so far been limited by acquired drug resistance. GB cells show a high mutation rate and usually present more than two sub-clones within the same patient and from the same primary tumour (McGranahan and Swanton, 2017; Qazi et al., 2017). The current literature suggest that a multiple approach is required to obtain better results (Prasad et al., 2011; Taylor et al., 2012; Westhoff et al., 2014; Westphal et al., 2017; Zhao et al., 2017), and we propose that GB and the surrounding healthy tissue communication is a central mediator of tumour progression.

### Progressive tumour growth

Although there is an overall significant expansion in volume of the glial TMs membrane and an increase in the number of glial cells in the *Drosophila* model of GB, there is no early significant increase in those parameters. Tumour volume shows some values above the average 2 days after tumour induction, and a great dispersion in the volume of the network and the number of glial cells on day 3 after tumour induction. We propose that under physiological conditions, there is a stable state that prevents cells from exiting the cell cycle, dividing or activating the actin cytoskeleton to extend a network of membrane projections. Therefore, on day 1 after tumour induction, GB cells do not show evident changes, there is a uniform range of values for the volume of membrane projections and number of glial cells. On day 2 after tumour induction, some individuals abandon this stable state and GB cells extend larger and longer membrane projections, although most individuals continue in stability and display morphological characteristics similar to control samples. However, on day 3 after tumour induction there is a great expansion of membrane projections and the number of glial cells is increased. Similar to GB patients, some individuals are more resistant than others to the loss of homeostasis, which would explain the variance in the values for network volume and number of glial cells on day 3 after tumour induction.

### Progressive neurodegeneration

There is a significant progressive decrease in the number of synapses at the NMJ. Nevertheless, the variance in the number of synapses on day 1 after tumour induction is very large, reaching uniform values on days 2 and 3 after tumour induction. The individuals show different resistances to the changes caused by the GB. On day 1 after tumour induction, some individuals are largely affected by the tumour and suffer a great loss of synapses at the NMJ while other individuals are more resistant, and the number of synapses remains normal. On day 2 after tumour induction, the impact of GB causes severe changes to all individuals regardless of their initial resistance.

We have observed a negative correlation between the volume of the membrane network and the number of synapses at the NMJ, suggesting that the expansion of the membrane projections is responsible for the neurodegeneration. Previous studies from our laboratory have proved that TMs surround neurons and deplete WNT from the neurons (Portela et al., 2019), which leads to synapse loss and neurodegeneration (Rich and Bigner, 2004). We can conclude that as the tumor progresses, TMs grow progressively, infiltrate in the brain and surround neurons leading to their degeneration, process previously described as vampirization (Portela et al., 2019). The three main events described here do not occur concomitantly, the progressive neurodegeneration is an early event in GB development, associated to early neuronal Wg/WNT depletion by GB cells. The expansion of the membrane network of TMs is slow during the first 24h of tumor development but increases after 48h of tumor development. TMs expansion and infiltration require Wg/WNT and JNK/MMPs pathways activation. Consequently, TMs expansion facilitate GB cells to further contact and communicate with the surrounding tissue. Eventually, Wg/WNT and JNK signals promote the increase in the number of GB cells that occurs as a later event. These evidences indicate that GB-neuron interaction is an early event required for GB progression. Consistent with this, a recent study in *Drosophila* EGFR-driven epithelial tumor model showed inter-tissue communication between epithelial and the adjacent healthy mesenchymal cells. The tumoral epithelial cells upregulate and transport the Notch ligand Delta in cytonemes, that contact and communicate with the healthy mesenchymal cells to activate Notch signaling in a non-autonomous manner. Notch activity keep the mesenchymal cell precursors in an undifferentiated and proliferative state that is critical to sustain tumor progression (Boukhatmi et al., 2020). This cell to cell interaction is similar to our findings in glia– neuron communication in GB, where glial cells use cytonemes/TMs to hijack neuronal Egr and activate JNK signaling in the tumor cells to sustain tumor progression.

### Progressive activation of JNK pathway via Egr/Grnd

One of the main molecular events in GB is the activation of JNK signalling that regulates MMPs expression, TMs expansion and infiltration. Previous results indicate that Wg pathway responds to JNK and network expansion, these three events were described as a regulatory positive feedback loop in GB progression. However, the signals that specifically activate JNK pathway are under debate. Grnd is the receptor that upon binding to the ligand Egr, activates JNK pathway. Our qPCR results and the localization analysis of Egr-GFP fused protein showed that Egr is produced in the surrounding neurons, and re-localized to the membrane of GB projections in contact with healthy neuronal tissue. The progressive activation of JNK pathway in glial cells correlate with the morphological changes (membrane network expansion and number of glial cells) that GB undergo 48h after tumor induction. Therefore, this example of neuron-GB interaction where non-autonomous signals facilitate tumor progression and GB-mediated neurodegeneration, contribute to the complexity and versatility of these incurable tumours. This neuron-GB communication mechanism highlights the relevance of the interactions and signalling between GB and the surrounding healthy tissue. Therefore, it is of interest to uncover the regulatory mechanisms that mediate *Egr* expression and secretion in neurons in response to GB induction, and the associated response in GB cells mediated by Grnd/Egr as a potential modulator of GB progression.

## Acknowledgements

We thank Professor Alberto Ferrús for helpful discussions. We are grateful to R. Read, I. Guerrero, P. Leopold,, E. Martín-Blanco, E. Moreno, the Bloomington *Drosophila* stock Centre and the Developmental Studies Hydridoma Bank for supplying fly stocks and antibodies, and FlyBase for its wealth of information. We acknowledge the support of the Confocal Microscopy unit and Molecular Biology unit at the Cajal Institute for their help with this project. MP holds a fellowship from the Juan de la Cierva program IJCI-2014- 19272 and SCT holds a contract from the Ramón y Cajal program RYC-2012-11410 from the Spanish MICINN. Research has been funded by grant BFU2015-65685P. Authors declare no conflicts of interest.

## Experimental Procedures

### Fly stocks

Flies were raised in standard fly food at 25°C. Fly stocks from the Bloomington stock Centre: *UAS-lacZ* (BL8529), *UAS-myr-RFP* (BL7119), *repo-Gal4* (BL7415), *tub-gal80*^*ts*^ (BL7019), *Egr-GFP* (BL66381), *UAS-egr-RNAi* (BL 55276 and BL58993) *UAS-dEGFR*^*λ*^, *UAS-PI3K92E* (*dp110*^*CAAX*^) (A gift from R. Read), *UAS-ihog-RFP* (a gift from I. Guerrero), *puc-lacZ* (a gift from E. Martín-Blanco), *UAS-dmyc (Krengel et al., 2004), UAS-TOR- DER*^*CA*^*(Dominguez et al., 1998)*.

### *Drosophila* glioblastoma model

In *Drosophila*, a combination of EGFR and PI3K mutations effectively causes a glioma-like condition that shows features of human gliomas including glia expansion, brain invasion, neuron dysfunction, synapse loss and neurodegeneration (Kegelman et al., 2014; Portela et al., 2019; Read, 2011; Read et al., 2009). To generate a glioma in *Drosophila* melanogaster, the Gal4/UAS system (Brand and Perrimon, 1993) was used as described above (*repo*-Gal4>UAS-*EGFR*^*λ*^,UAS-*dp110*^*CAAX*^. To restrict the expression of this genetic combination and control it in a temporal manner, we used the thermo sensitive repression system Gal80^TS^. Individuals maintained at 17°C did not activate the expression of the UAS constructs, but when the larvae were switched to 29°C, the protein Gal80^TS^ changed conformation and was not longer able to bind to Gal4 to prevent its interaction with UAS sequences, and the expression system was activated and therefore the GB was induced. *Tub-GAL80*^*TS*^; *repo*-Gal4>UAS-*EGFR*^*λ*^, UAS-*dp110*^*CAAX*^ animals were raised at 17 C, shifted to 29 C for 1, 2, 3 or 4 days at 8, 6.3, 3.7 or 2 days after egg laying (AEL) respectively, and subjected to dissection immediately after.

### Antibodies for Immunofluorescence

Third-instar larval brains, were dissected in phosphate-buffered saline (PBS), fixed in 4% formaldehyde for 30min, washed in PBS + 0.1 or 0.3% Triton X-100 (PBT), and blocked in PBT + 5% BSA.

Antibodies used were: mouse anti-Wg (DSHB 1:50), mouse anti-Repo (DSHB 1:50), mouse anti-Fz1 (DSHB 1:50), mouse anti-Cyt-Arm (DSHB 1:50), mouse anti-MMP1 (DSHB 5H7B11, 3A6B4, 3B8D12, 1:50), guinea pig anti-Grnd (Andersen et al., 2015) (1:250, P. Leopold), mouse anti-β-galactosidase (Sigma, 1:500), mouse anti-GFP (Invitrogen A11120, 1:500), mouse anti-bruchpilot (DSHB Nc82, 1:20), Rabbit anti-Hrp (Jackson Immunoresearch 111-035-144, 1:400).

Secondary antibodies: anti-mouse Alexa 488, 568, 647, anti-rabbit Alexa 488, 568, 647 (Thermofisher, 1:500). DNA was stained with 2-(4-amidinophenyl)-1H-indole-6- carboxamidine (DAPI, 1µM).

### Imaging

Fluorescent labelled samples were mounted in Vectashield mounting media with DAPI (Vector Laboratories) and analysed by Confocal microscopy (LEICA TCS SP5). In all experiments the whole brain lobes were acquired individually and whole stacks were analysed. The images shown in the figures are a single plane images to facilitate visualization. In some panels, higher magnifications of single plane images are shown in the figures to facilitate the visualization of the antibody staining’s in more detail. Images were processed using Leica LAS AF Lite and Fiji (Image J 1.50e). Images were assembled using Adobe Photoshop CS5.1.

### Quantifications

Relative MMP1 and Grnd staining within brains was determined from images taken at the same confocal settings. Average pixel intensity was measured using measurement log tool from Fiji 1.51g and Adobe Photoshop CS5.1. Average pixel intensity was measured in the glial tissue and in the adjacent neuronal tissue (N<10 for each sample) and expressed as a Glia/Neuron ratio. Glial network volume was quantified using Imaris surface tool (Imaris 6.3.1 Bitplane Scientific Solutions software).

The number of Repo^+^ cells, the number of synaptic active sites and the number of puc-lacZ positive cells was quantified by using the spots tool Imaris 6.3.1 software, we selected a minimum size and threshold for the puncta in the control samples of each experiment. Then we applied these conditions to the analysis of each corresponding experimental sample. For the puc-lacZ glia or neuron co-localization studies we quantified the total number of puc-lacZ^+^ cells and then applied a co-localization filter (intensity center of the channel of interest) using the Spots tool from the Imaris 6.3.1 software.

For the co-localization of Egr-GFP in glial cells, GFP channel volume was quantified using Imaris surface tool. We selected a specific threshold for the total volume in the control samples and then we applied these conditions to the analysis of the corresponding experimental sample. Then we applied a co-localization filter (intensity mean of the red channel)

## Statistical Analysis

To analyse and plot the data, we used Microsoft Excel 2013 and GraphPad Prism 6. We performed a D’Agostino & Pearson normality test and the data found to have a normal distribution were analysed by a two-tailed t test with Welch-correction. In the case of multiple comparisons, we used a One-way ANOVA with Bonferroni post-test. The data that did not pass the normality test were subjected to a two-tailed Mann–Whitney U test or in the case of multiple comparisons a Kruskal–Wallis test with Dunns post-test. Error bars represent standard error of the mean. * represents p value ≤ 0.05; ** p value ≤ 0.01; *** p value ≤ 0.001. Statistical values of p value >0.05 were not considered significant, (n.s.).

## Figure Legends

**Figure S1:**
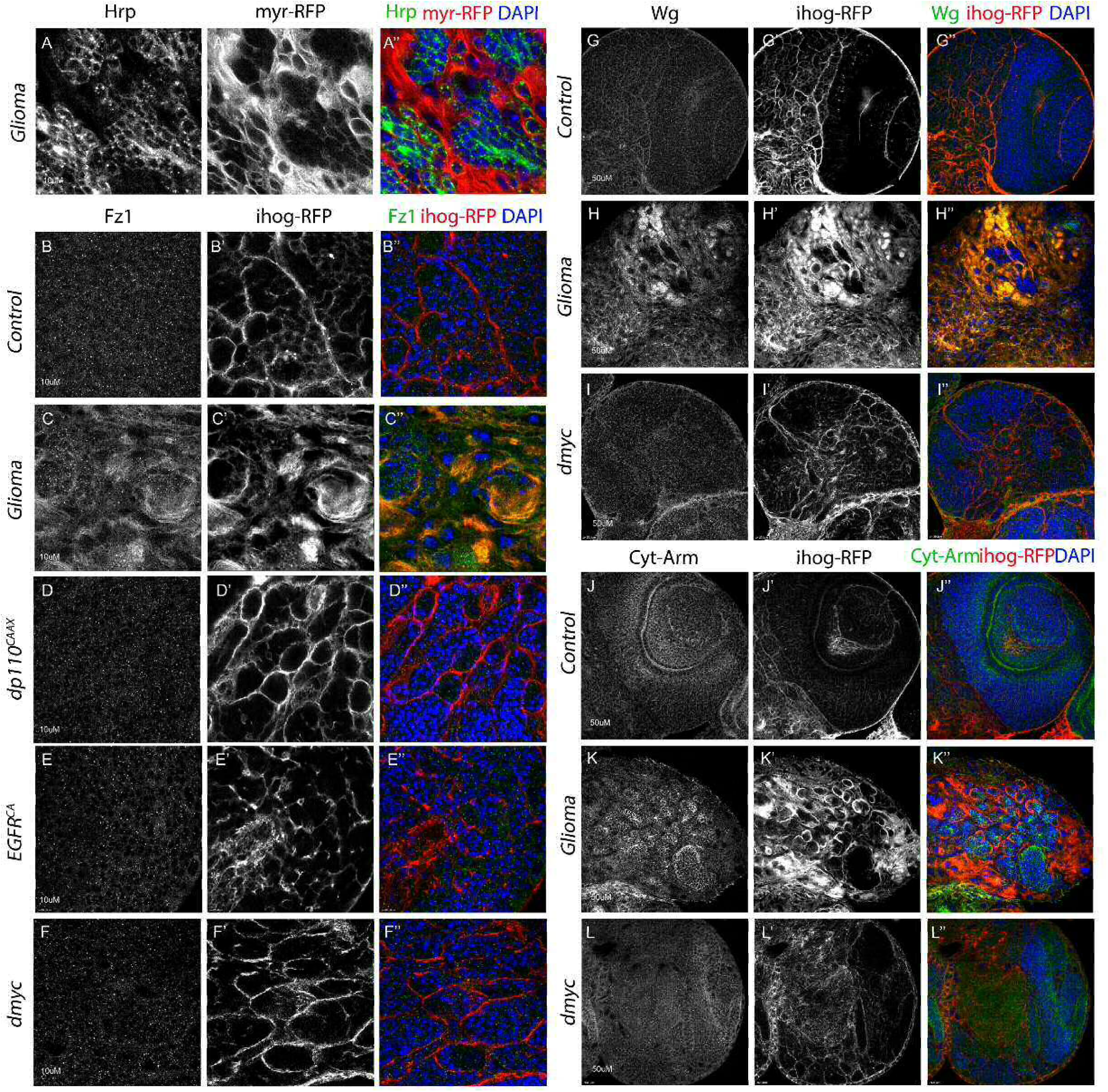
Independent constitutive activation of PI3K or EGFR or ectopic *dmyc* are not responsible for Wg/Fz1 accumulation and expansion of TM network. Brains from 3rd instar larvae. (A) Glia is labeled with *UAS-myr-RFP* driven by *repo-Gal4* to visualize membrane projections in glial cells (red) and neurons are stained with Hrp (green) and enwrapped by glial TMs in glioma brains (merge). (B-F) Glia are labeled with *UAS-Ihog-RFP* driven by *repo-Gal4* to visualize active cytonemes/ TM structures in glial cells and stained with Fz1/Wg or Cyt-Arm (green). (B-F) Fz1 (green) is homogeneously distributed in control (B). (C) Fz1 accumulates in the TMs in glioma brains. Fz1 is homogeneously distributed in (D) *dp110*^*CAAX*^, (E)*EGFR*^*CA*^ and (F) *dmyc* sections similar to the control (B). (G-L) Wg and Cyt-Arm (green) are homogeneously distributed in control brain sections (G, J) as well as in *dmyc* sections (I, L) and the glial network shown in red by Ihog-RFP does not overgrow or encapsulate neuronal clusters as opposed to GB brains where Wg accumulates in TMs (H) and Cyt-Arm accumulates in the neurons’ cytoplasm where it is inactive (K). Nuclei are marked with DAPI. Genotypes: (A) *UAS-dEGFR*^*λ*^, *UAS-dp110*^*CAAX*^; *Gal80*^*ts*^; *repo-Gal4, myr-RFP* (B, G, J) *UAS-lacZ/repo-Gal4, UAS-ihog-RFP* (C, H, K) *UAS-dEGFR*^*λ*^, *UAS-dp110*^*CAAX*^;; *repo-Gal4, UAS-ihog-RFP* (D) *UAS-dp110*^*CAAX*^;; *repo-Gal4, UAS-ihog-RFP* (E) *UAS-TOR-DER*^*CA*^;; *repo-Gal4, UAS-ihog-RFP* (F, I, L) *repo-Gal4, UAS-ihog-RFP/UAS-dmyc*

## References

Alifieris, C. and Trafalis, D. T. (2015). Glioblastoma multiforme: Pathogenesis and treatment. Pharmacol Ther 152, 63–82.

Andersen, D. S., Colombani, J., Palmerini, V., Chakrabandhu, K., Boone, E., Rothlisberger, M., Toggweiler, J., Basler, K., Mapelli, M., Hueber, A. O. et al. (2015). The Drosophila TNF receptor Grindelwald couples loss of cell polarity and neoplastic growth. Nature 522, 482–6.

Annibali, D., Whitfield, J. R., Favuzzi, E., Jauset, T., Serrano, E., Cuartas, I., Redondo-Campos, S., Folch, G., Gonzalez-Junca, A., Sodir, N. M. et al. (2014). Myc inhibition is effective against glioma and reveals a role for Myc in proficient mitosis. Nat Commun 5, 4632.

Arnes, M. and Casas Tinto, S. (2017). Aberrant Wnt signaling: a special focus in CNS diseases. J Neurogenet 31, 216–222.

Boukhatmi, H., Martins, T., Pillidge, Z., Kamenova, T. and Bray, S. (2020). Notch Mediates Inter-tissue Communication to Promote Tumorigenesis. Curr Biol.

Brand, A. H. and Perrimon, N. (1993). Targeted gene expression as a means of altering cell fates and generating dominant phenotypes. Development 118, 401–15.

Casas-Tinto, S., Arnes, M. and Ferrus, A. (2017). Drosophila enhancer-Gal4 lines show ectopic expression during development. R Soc Open Sci 4, 170039.

Casas-Tinto, S. and Portela, M. (2019). Cytonemes, Their Formation, Regulation, and Roles in Signaling and Communication in Tumorigenesis. Int J Mol Sci 20.

Chatterjee, N. and Bohmann, D. (2012). A versatile PhiC31 based reporter system for measuring AP-1 and Nrf2 signaling in Drosophila and in tissue culture. PLoS One 7, e34063.

Chen, W., Zhong, X., Wei, Y., Liu, Y., Yi, Q., Zhang, G., He, L., Chen, F. and Luo, J. (2016). TGF-beta Regulates Survivin to Affect Cell Cycle and the Expression of EGFR and MMP9 in Glioblastoma. Mol Neurobiol 53, 1648–1653.

Cheng, C. Y., Hsieh, H. L., Hsiao, L. D. and Yang, C. M. (2012). PI3-K/Akt/JNK/NF-kappaB is essential for MMP-9 expression and outgrowth in human limbal epithelial cells on intact amniotic membrane. Stem Cell Res 9, 9–23.

Cherry, E. M., Lee, D. W., Jung, J. U. and Sitcheran, R. (2015). Tumor necrosis factor-like weak inducer of apoptosis (TWEAK) promotes glioma cell invasion through induction of NF-kappaB-inducing kinase (NIK) and noncanonical NF-kappaB signaling. Mol Cancer 14, 9.

Chicheportiche, Y., Bourdon, P. R., Xu, H., Hsu, Y. M., Scott, H., Hession, C., Garcia, I. and Browning, J. L. (1997). TWEAK, a new secreted ligand in the tumor necrosis factor family that weakly induces apoptosis. J Biol Chem 272, 32401–10.

de Lucas, A. G., Schuhmacher, A. J., Oteo, M., Romero, E., Camara, J. A., de Martino, A., Arroyo, A. G., Morcillo, M. A., Squatrito, M., Martinez-Torrecuadrada, J. L. et al. (2016). Targeting MT1-MMP as an ImmunoPET-Based Strategy for Imaging Gliomas. PLoS One 11, e0158634.

Dominguez, M., Wasserman, J. D. and Freeman, M. (1998). Multiple functions of the EGF receptor in Drosophila eye development. Curr Biol 8, 1039–48.

Feng, J., Yan, P. F., Zhao, H. Y., Zhang, F. C., Zhao, W. H. and Feng, M. (2016). Inhibitor of Nicotinamide Phosphoribosyltransferase Sensitizes Glioblastoma Cells to Temozolomide via Activating ROS/JNK Signaling Pathway. Biomed Res Int 2016, 1450843.

Furnari, F. B., Fenton, T., Bachoo, R. M., Mukasa, A., Stommel, J. M., Stegh, A., Hahn, W. C., Ligon, K. L., Louis, D. N., Brennan, C. et al. (2007). Malignant astrocytic glioma: genetics, biology, and paths to treatment. Genes Dev 21, 2683–710.

Gallego, O. (2015). Nonsurgical treatment of recurrent glioblastoma. Curr Oncol 22, e273–81.

Hagemann, C., Anacker, J., Ernestus, R. I. and Vince, G. H. (2012). A complete compilation of matrix metalloproteinase expression in human malignant gliomas. World J Clin Oncol 3, 67–79.

Hagemann, C., Anacker, J., Haas, S., Riesner, D., Schomig, B., Ernestus, R. I. and Vince, G. H. (2010). Comparative expression pattern of Matrix-Metalloproteinases in human glioblastoma cell-lines and primary cultures. BMC Res Notes 3, 293.

Hagemann, T., Wilson, J., Kulbe, H., Li, N. F., Leinster, D. A., Charles, K., Klemm, F., Pukrop, T., Binder, C. and Balkwill, F. R. (2005). Macrophages induce invasiveness of epithelial cancer cells via NF-kappa B and JNK. J Immunol 175, 1197–205.

Hayden, E. C. (2010). Genomics boosts brain-cancer work. Nature 463, 278.

Holland, E. C. (2000). Glioblastoma multiforme: the terminator. Proc Natl Acad Sci U S A 97, 6242–4.

Hsu, J. B., Chang, T. H., Lee, G. A., Lee, T. Y. and Chen, C. Y. (2019). Identification of potential biomarkers related to glioma survival by gene expression profile analysis. BMC Med Genomics 11, 34.

Huang, C., Rajfur, Z., Borchers, C., Schaller, M. D. and Jacobson, K. (2003). JNK phosphorylates paxillin and regulates cell migration. Nature 424, 219–23.

Igaki, T., Kanda, H., Yamamoto-Goto, Y., Kanuka, H., Kuranaga, E., Aigaki, T. and Miura, M. (2002). Eiger, a TNF superfamily ligand that triggers the Drosophila JNK pathway. EMBO J 21, 3009–18.

Ispanovic, E. and Haas, T. L. (2006). JNK and PI3K differentially regulate MMP-2 and MT1-MMP mRNA and protein in response to actin cytoskeleton reorganization in endothelial cells. Am J Physiol Cell Physiol 291, C579–88.

Jarabo, P., de Pablo, C., Herranz, H., Martin, F. A. and Casas-Tinto, S. (2020).

Jemc, J. C., Milutinovich, A. B., Weyers, J. J., Takeda, Y. and Van Doren, M. (2012). raw Functions through JNK signaling and cadherin-based adhesion to regulate Drosophila gonad morphogenesis. Dev Biol 367, 114–25.

Kamino, M., Kishida, M., Kibe, T., Ikoma, K., Iijima, M., Hirano, H., Tokudome, M., Chen, L., Koriyama, C., Yamada, K. et al. (2011). Wnt-5a signaling is correlated with infiltrative activity in human glioma by inducing cellular migration and MMP-2. Cancer Sci 102, 540–8.

Kegelman, T. P., Hu, B., Emdad, L., Das, S. K., Sarkar, D. and Fisher, P. B. (2014). In vivo modeling of malignant glioma: the road to effective therapy. Adv Cancer Res 121, 261–330.

Keshishian, H., Broadie, K., Chiba, A. and Bate, M. (1996). The drosophila neuromuscular junction: a model system for studying synaptic development and function. Annu Rev Neurosci 19, 545–75.

Kitanaka, C., Sato, A. and Okada, M. (2013). JNK Signaling in the Control of the Tumor-Initiating Capacity Associated with Cancer Stem Cells. Genes Cancer 4, 388–96.

Krengel, U., Olsson, L. L., Martinez, C., Talavera, A., Rojas, G., Mier, E., Angstrom, J. and Moreno, E. (2004). Structure and molecular interactions of a unique antitumor antibody specific for N-glycolyl GM3. J Biol Chem 279, 5597–603.

LaFever, K. S., Wang, X., Page-McCaw, P., Bhave, G. and Page-McCaw, A. (2017). Both Drosophila matrix metalloproteinases have released and membrane-tethered forms but have different substrates. Sci Rep 7, 44560.

Langen, M., Koch, M., Yan, J., De Geest, N., Erfurth, M. L., Pfeiffer, B. D., Schmucker, D., Moreau, Y. and Hassan, B. A. (2013). Mutual inhibition among postmitotic neurons regulates robustness of brain wiring in Drosophila. Elife 2, e00337.

Lee, Y. S., Lan Tran, H. T. and Van Ta, Q. (2009). Regulation of expression of matrix metalloproteinase-9 by JNK in Raw 264.7 cells: presence of inhibitory factor(s) suppressing MMP-9 induction in serum and conditioned media. Exp Mol Med 41, 259–68.

Louis, D. N., Perry, A., Reifenberger, G., von Deimling, A., Figarella-Branger, D., Cavenee, W. K., Ohgaki, H., Wiestler, O. D., Kleihues, P. and Ellison, D. W. (2016). The 2016 World Health Organization Classification of Tumors of the Central Nervous System: a summary. Acta Neuropathol 131, 803–20.

Lowy, A. M., Clements, W. M., Bishop, J., Kong, L., Bonney, T., Sisco, K., Aronow, B., Fenoglio-Preiser, C. and Groden, J. (2006). beta-Catenin/Wnt signaling regulates expression of the membrane type 3 matrix metalloproteinase in gastric cancer. Cancer Res 66, 4734–41.

Lyu, J. and Joo, C. K. (2005). Wnt-7a up-regulates matrix metalloproteinase-12 expression and promotes cell proliferation in corneal epithelial cells during wound healing. J Biol Chem 280, 21653–60.

Maher, E. A., Furnari, F. B., Bachoo, R. M., Rowitch, D. H., Louis, D. N., Cavenee, W. K. and DePinho, R. A. (2001). Malignant glioma: genetics and biology of a grave matter. Genes Dev 15, 1311–33.

Malemud, C. J. (2006). Matrix metalloproteinases (MMPs) in health and disease: an overview. Front Biosci 11, 1696–701.

Martin-Blanco, E., Gampel, A., Ring, J., Virdee, K., Kirov, N., Tolkovsky, A. M. and Martinez-Arias, A. (1998). puckered encodes a phosphatase that mediates a feedback loop regulating JNK activity during dorsal closure in Drosophila. Genes Dev 12, 557–70.

Matsuda, K., Sato, A., Okada, M., Shibuya, K., Seino, S., Suzuki, K., Watanabe, E., Narita, Y., Shibui, S., Kayama, T. et al. (2012). Targeting JNK for therapeutic depletion of stem-like glioblastoma cells. Sci Rep 2, 516.

McGranahan, N. and Swanton, C. (2017). Clonal Heterogeneity and Tumor Evolution: Past, Present, and the Future. Cell 168, 613–628.

McGuire, S. (2016). World Cancer Report 2014. Geneva, Switzerland: World Health Organization, International Agency for Research on Cancer, WHO Press, 2015. Adv Nutr 7, 418–9.

Miller, J. J., Shih, H. A., Andronesi, O. C. and Cahill, D. P. (2017). Isocitrate dehydrogenase-mutant glioma: Evolving clinical and therapeutic implications. Cancer 123, 4535–4546.

Moreno, E., Yan, M. and Basler, K. (2002). Evolution of TNF signaling mechanisms: JNK-dependent apoptosis triggered by Eiger, the Drosophila homolog of the TNF superfamily. Curr Biol 12, 1263–8.

Mu, N., Gu, J., Liu, N., Xue, X., Shu, Z., Zhang, K., Huang, T., Chu, C., Zhang, W., Gong, L. et al. (2018). PRL-3 is a potential glioblastoma prognostic marker and promotes glioblastoma progression by enhancing MMP7 through the ERK and JNK pathways. Theranostics 8, 1527–1539.

Munaut, C., Noel, A., Hougrand, O., Foidart, J. M., Boniver, J. and Deprez, M. (2003). Vascular endothelial growth factor expression correlates with matrix metalloproteinases MT1-MMP, MMP-2 and MMP-9 in human glioblastomas. Int J Cancer 106, 848–55.

Nakada, M., Okada, Y. and Yamashita, J. (2003). The role of matrix metalloproteinases in glioma invasion. Front Biosci 8, e261–9.

Okada, M., Sato, A., Shibuya, K., Watanabe, E., Seino, S., Suzuki, S., Seino, M., Narita, Y., Shibui, S., Kayama, T. et al. (2014). JNK contributes to temozolomide resistance of stem-like glioblastoma cells via regulation of MGMT expression. Int J Oncol 44, 591–9.

Osswald, M., Jung, E., Sahm, F., Solecki, G., Venkataramani, V., Blaes, J., Weil, S., Horstmann, H., Wiestler, B., Syed, M. et al. (2015). Brain tumour cells interconnect to a functional and resistant network. Nature 528, 93–8.

Osswald, M., Solecki, G., Wick, W. and Winkler, F. (2016). A malignant cellular network in gliomas: potential clinical implications. Neuro Oncol 18, 479–85.

Page-McCaw, A., Serano, J., Sante, J. M. and Rubin, G. M. (2003). Drosophila matrix metalloproteinases are required for tissue remodeling, but not embryonic development. Dev Cell 4, 95–106.

Portela, M., Venkataramani, V., Fahey-Lozano, N., Seco, E., Losada-Perez, M., Winkler, F. and Casas-Tinto, S. (2019). Glioblastoma cells vampirize WNT from neurons and trigger a JNK/MMP signaling loop that enhances glioblastoma progression and neurodegeneration. PLoS Biol 17, e3000545.

Prasad, G., Sottero, T., Yang, X., Mueller, S., James, C. D., Weiss, W. A., Polley, M. Y., Ozawa, T., Berger, M. S., Aftab, D. T. et al. (2011). Inhibition of PI3K/mTOR pathways in glioblastoma and implications for combination therapy with temozolomide. Neuro Oncol 13, 384–92.

Qazi, M. A., Vora, P., Venugopal, C., Sidhu, S. S., Moffat, J., Swanton, C. and Singh, S. K. (2017). Intratumoral heterogeneity: pathways to treatment resistance and relapse in human glioblastoma. Ann Oncol 28, 1448–1456.

Read, R. D. (2011). Drosophila melanogaster as a model system for human brain cancers. Glia 59, 1364–76.

Read, R. D., Cavenee, W. K., Furnari, F. B. and Thomas, J. B. (2009). A drosophila model for EGFR-Ras and PI3K-dependent human glioma. PLoS Genet 5, e1000374.

Rich, J. N. and Bigner, D. D. (2004). Development of novel targeted therapies in the treatment of malignant glioma. Nat Rev Drug Discov 3, 430–46.

Rogers, T. W., Toor, G., Drummond, K., Love, C., Field, K., Asher, R., Tsui, A., Buckland, M. and Gonzales, M. (2018). The 2016 revision of the WHO Classification of Central Nervous System Tumours: retrospective application to a cohort of diffuse gliomas. J Neurooncol 137, 181–189.

Roomi, M. W., Kalinovsky, T., Rath, M. and Niedzwiecki, A. (2017). Modulation of MMP-2 and MMP-9 secretion by cytokines, inducers and inhibitors in human glioblastoma T-98G cells. Oncol Rep 37, 1907–1913.

Roth, W., Wild-Bode, C., Platten, M., Grimmel, C., Melkonyan, H. S., Dichgans, J. and Weller, M. (2000). Secreted Frizzled-related proteins inhibit motility and promote growth of human malignant glioma cells. Oncogene 19, 4210–20.

Ruan, W., Srinivasan, A., Lin, S., Kara k, I. and Barker, P. A. (2016). Eiger-induced cell death relies on Rac1-dependent endocytosis. Cell Death Dis 7, e2181.

Tateishi, K., Iafrate, A. J., Ho, Q., Curry, W. T., Batchelor, T. T., Flaherty, K. T., Onozato, M. L., Lelic, N., Sundaram, S., Cahill, D. P. et al. (2016). Myc-Driven Glycolysis Is a Therapeutic Target in Glioblastoma. Clin Cancer Res 22, 4452–65.

Taylor, T. E., Furnari, F. B. and Cavenee, W. K. (2012). Targeting EGFR for treatment of glioblastoma: molecular basis to overcome resistance. Curr Cancer Drug Targets 12, 197–209.

Uhlirova, M. and Bohmann, D. (2006). JNK- and Fos-regulated Mmp1 expression cooperates with Ras to induce invasive tumors in Drosophila. EMBO J 25, 5294–304.

Uraguchi, M., Morikawa, M., Shirakawa, M., Sanada, K. and Imai, K. (2004). Activation of WNT family expression and signaling in squamous cell carcinomas of the oral cavity. J Dent Res 83, 327–32.

Veeravalli, K. K. and Rao, J. S. (2012). MMP-9 and uPAR regulated glioma cell migration. Cell Adh Migr 6, 509–12.

Waitkus, M. S., Diplas, B. H. and Yan, H. (2018). Biological Role and Therapeutic Potential of IDH Mutations in Cancer. Cancer Cell 34, 186–195.

Wang, X., Huang, Z., Wu, Q., Prager, B. C., Mack, S. C., Yang, K., Kim, L. J. Y., Gimple, R. C., Shi, Y., Lai, S. et al. (2017). MYC-Regulated Mevalonate Metabolism Maintains Brain Tumor-Initiating Cells. Cancer Res 77, 4947–4960.

Westhoff, M. A., Karpel-Massler, G., Bruhl, O., Enzenmuller, S., La Ferla-Bruhl, K., Siegelin, M. D., Nonnenmacher, L. and Debatin, K. M. (2014). A critical evaluation of PI3K inhibition in Glioblastoma and Neuroblastoma therapy. Mol Cell Ther 2, 32.

Westphal, M., Maire, C. L. and Lamszus, K. (2017). EGFR as a Target for Glioblastoma Treatment: An Unfulfilled Promise. CNS Drugs 31, 723–735.

Wick, W., Osswald, M., Wick, A. and Winkler, F. (2018). Treatment of glioblastoma in adults. Ther Adv Neurol Disord 11, 1756286418790452.

Yamamoto, M., Ueno, Y., Hayashi, S. and Fukushima, T. (2002). The role of proteolysis in tumor invasiveness in glioblastoma and metastatic brain tumors. Anticancer Res 22, 4265–8.

Zeigler, M. E., Chi, Y., Schmidt, T. and Varani, J. (1999). Role of ERK and JNK pathways in regulating cell motility and matrix metalloproteinase 9 production in growth factor-stimulated human epidermal keratinocytes. J Cell Physiol 180, 271–84.

Zeng, A., Yin, J., Li, Y., Li, R., Wang, Z., Zhou, X., Jin, X., Shen, F., Yan, W. and You, Y. (2018). miR-129-5p targets Wnt5a to block PKC/ERK/NF-kappaB and JNK pathways in glioblastoma. Cell Death Dis 9, 394.

Zhao, H. F., Wang, J., Shao, W., Wu, C. P., Chen, Z. P., To, S. T. and Li, W. P. (2017). Recent advances in the use of PI3K inhibitors for glioblastoma multiforme: current preclinical and clinical development. Mol Cancer 16, 100.

